# Road proximity differentially shapes rodent-mediated seed dispersal frequency and distance

**DOI:** 10.64898/2026.03.24.713877

**Authors:** João Craveiro, Miguel N. Bugalho, Pedro G. Vaz

## Abstract

By concentrating rodents along verges, roads can reshape rodent-mediated seed dispersal, yet empirical tests remain scarce. We conducted a two-year field experiment in Mediterranean oak woodlands in southern Portugal to test how seed dispersal varies with distance from roads across road type (paved vs. unpaved) and road–forest context (edge vs. non-edge). We tracked labeled holm oak acorns, recording dispersal distances and the number of dispersal events. The two metrics responded differently to road distance. Dispersal distances changed little with distance from roads in non-edge contexts but increased in edge road–forest contexts (2× longer at 400 m than at 10 m) and showed a year × distance-to-road interaction, with longer dispersal distances farther from roads in the second year (a poor mast year). Dispersal distances were also longer when acorns were deposited under shrubs and in areas of higher tree density, and decreased with greater natural acorn availability. In contrast, the number of dispersal events declined with distance from roads (30% more events at 10 m than at 400 m) and was higher along unpaved than paved roads (39% more events). Dispersal frequency also increased in the poor mast year and with shrub cover. No acorns crossed the road. Thus, road verges can concentrate rodent seed handling but do not increase dispersal distances near roads nor provide cross-road seed connectivity; instead, dispersal outcomes depend on edge context, road type, and microhabitat structure. Management that retains structural cover at verges and the adjacent forest edge (e.g., shrub patches and non-uniform clearing) can harness verge-associated activity to increase acorn deposition in sheltered microsites and promote regeneration farther into forest interiors in roaded landscapes.

## INTRODUCTION

Roads exert multiple impacts on habitats and wildlife, disrupting key ecological processes such as plant-animal interactions by altering animal movement and landscape connectivity (Craveiro et al., 2019; Cui et al., 2018; Markl et al., 2012; Quiles & Barrientos, 2024). One example is the disruption of animal-mediated seed dispersal, a crucial ecological function sustaining forest regeneration (Chen et al., 2019a; Nathan & Muller-Landau, 2000; Wang & Smith, 2002). The spatial footprint of road impacts can penetrate variably into surrounding habitats—the “road-effect zone” (Chen et al., 2019b; Forman & Deblinger, 2000). Despite well-established evidence of broad ecological effects of roads (van der Ree et al., 2015), little is known about how roads affect animal-mediated seed dispersal across distance gradients—specifically the frequency of dispersal events and seed dispersal distances—and whether effects depend on road type and road–forest context.

Rodents, in particular, can be relatively tolerant of road-associated disturbance and occur at higher densities along verges than in forest interiors; nonetheless, they may avoid crossing road surfaces while carrying large seeds such as acorns, potentially constraining dispersal patterns (Fahrig & Rytwinski, 2009; Galantinho et al., 2017; Niu et al., 2021; Rosalino et al., 2011). Through scatter-hoarding—collecting and burying seeds in spatially dispersed caches—rodents promote germination and tree recruitment (Gómez et al., 2019; Perea et al., 2011a; Vaz et al., 2024). This behavior is especially critical in Iberian oak woodlands, where oak decline and poor regeneration are long-standing concerns (Bugalho et al., 2011; Vaz et al., 2019). Despite their ecological and economic value (Bugalho et al., 2009, 2011; Plieninger et al., 2011), these systems face multiple threats from climate change (Díaz et al., 2021), pathogens (Branco & Ramos, 2009; Brasier, 1992, 1996), phytophagous insects (Branco et al., 2002), wildfires (Moreira et al., 2011; Vaz et al., 2013), and overgrazing (Wadud et al., 2024). Understanding how rodent-mediated acorn dispersal operates at different distances from roads is thus crucial for developing effective strategies for oak woodland resilience.

Empirical results of how road proximity reshapes rodent-mediated dispersal along distance gradients remain scarce and mixed. Road proximity has been linked to reduced dispersal effectiveness and shorter dispersal distances that weaken with increasing distance into forest interiors (Chen et al., 2019b). Other studies report strong distance-dependent differences in dispersal effectiveness, yet persistently shorter dispersal distances near roads (Cui et al., 2018), as well as contrasting roadside–interior dynamics in removal rates and dispersal distances around paved roads (Niu et al., 2018, 2021). Together, this evidence suggests that road effects on seed dispersal frequency and distances may vary across contexts and road types.

Edges, as transitional zones between habitats, may modulate rodent foraging and dispersal dynamics (Coutant et al., 2022; Craveiro et al., 2025). In the highlands of Chiapas, Mexico, acorn removal by scatter-hoarding rodents (*Peromyscus* and *Reithrodontomys* spp.) was lower in open grasslands than in forested habitats, while dispersal activity was greater from forest edges toward interiors (López-Barrera et al., 2007). In northern China, distance from the forest edge influenced rodent seed removal rates, although dispersal distances did not differ significantly (Wang et al., 2025).

Road type may shape rodent-mediated dispersal by altering traffic intensity and substrate. In southern Portugal, rodents dispersed seeds farther along paved roadsides, primarily moving seeds parallel to roads, whereas crossings were infrequent and occurred more often along unpaved roads (Craveiro et al., 2025). Dispersal outcomes depend also on the microhabitat where acorns are deposited—such as open areas versus beneath shrubs—which can influence both dispersal frequency and dispersal distance (Evrard et al., 2017; Vaz et al., 2024). Overall, these studies suggest that distance-to-road effects on rodent-mediated dispersal may be shaped by road–forest context, road type, and microhabitat features, yet these factors are rarely tested simultaneously when quantifying dispersal frequency and dispersal distances.

We conducted a two-year field experiment in Mediterranean oak woodlands of southern Portugal to evaluate how a gradient of distance to road shapes rodent-mediated acorn dispersal, as well as the potential effects of road type (paved vs. unpaved) and road–forest context (edge vs. non-edge). Specifically, we asked: (i) How do dispersal distances and the number of dispersal events vary along a distance gradient to the road? We hypothesized that rodents would disperse acorns more often and over longer distances near the road with a gradual decrease with distance into the forest interior, reflecting rodent use of roadside verges as refuges. (ii) Do the effect of distance to road on dispersal patterns vary with road type and road–forest context? We hypothesized that rodents would disperse seeds more often and over longer distances in sites adjacent to unpaved roads, as the effects of paved roads are expected to extend farther into the forest interior than those of unpaved roads. Moreover, we expected higher dispersal rates in edge contexts than in non-edge contexts, where acorn availability can be lower. (iii) Does the effect of distance to the road vary interannually? We hypothesized that rodents would disperse seeds more often and over longer distances in a year with fewer acorns available.

## METHODS

### Study area and main study design

We conducted the study on 12 road sections (600 m each) in the Alentejo region, southern Portugal (Figure 1). The local climate is Mediterranean, with cold, wet winters and hot, dry summers (mean annual precipitation: 620 mm; mean temperature: 18 °C). The landscape consists of undulating terrain (200–400 m asl) dominated by montado (dehesa in Spain) and pastures. Montados are characterized by an open-canopy of cork (*Quercus suber*) and holm (*Quercus rotundifolia*) oak stands with an understory of shrubs (e.g., *Cistus*, *Ulex*, *Erica*, *Lavandula* spp.; Azedo et al., 2022; Bugalho et al., 2009, 2011; Castro & Freitas, 2009) and grassland areas. Grasslands in open areas consist mainly of annual forbs, graminoids, and legumes. Livestock, mainly cattle and sheep, with occasional domestic pigs, occurs in the region. Acorns naturally fall between September and January (Leiva & Perelló-Rodríguez, 2024). The main acorn-dispersing rodents in the area are the wood mouse (*Apodemus sylvaticus*) and the Algerian mouse (*Mus spretus*) (Galantinho et al., 2020).

**Figure 1.**
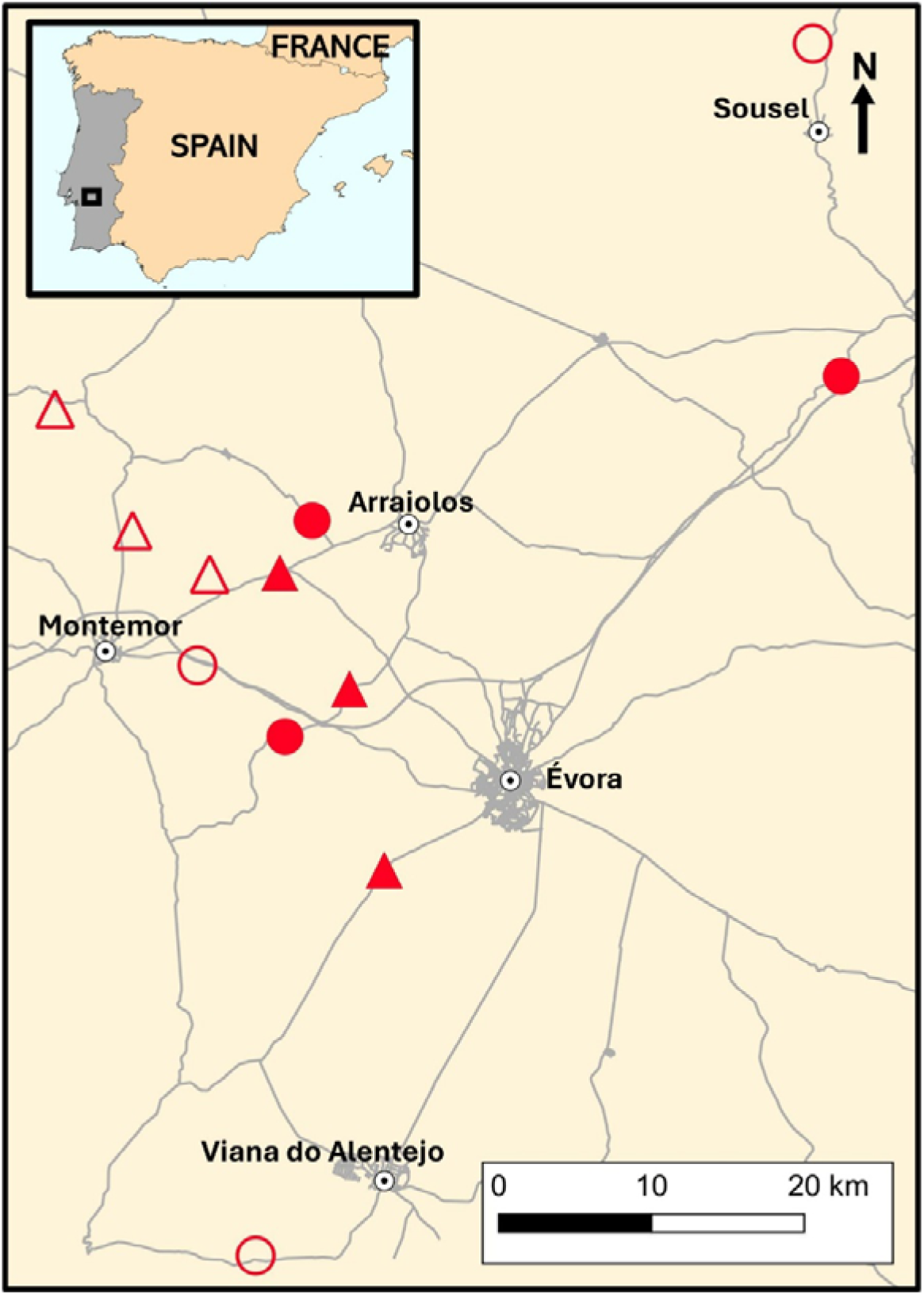
Location of the 12 road sections (600 m each) in the Alentejo region, southern Portugal, where seed dispersal by rodents was monitored. The experimental design included two treatments: (i) road-forest context (edge [circles] vs. non-edge [triangles]) and (ii) road type (paved [solid symbols] vs. unpaved [unfilled symbols]).

The 12 road sections (3.6–26.1 km apart; mean = 10.1 km) encompassed two road forest contexts—six bordering oak forests (edge) and six traversing oak forests (non-edge)—and two road types (six paved, six unpaved). Edge sites were characterized by oak woodland on one side and open pasture or cropland on the other. Because landscape directs animal movement, we chose edge and non-edge sites that were homogeneous in landscape composition. Paved roads (∼8 m wide; range: 7–11 m) were National Roads with asphalt pavement and a traffic volume of 112 vehicles/hour (range: 16–234), corresponding to 105 vehicles h□¹ (± 48 SE) on edge roads and 119 vehicles h□¹ (± 49 SE) on non-edge roads (see www.infraestruturasdeportugal.pt). Unpaved roads (∼4 m wide; range: 3–5 m) were public dirt roads with low traffic, mainly for agricultural access. To account for spatial variation, we established three replicates per factor (2 road-forest contexts × 2 road types × 3 replicates = 12 road sections) (Figure 2A).

**Figure 2.**
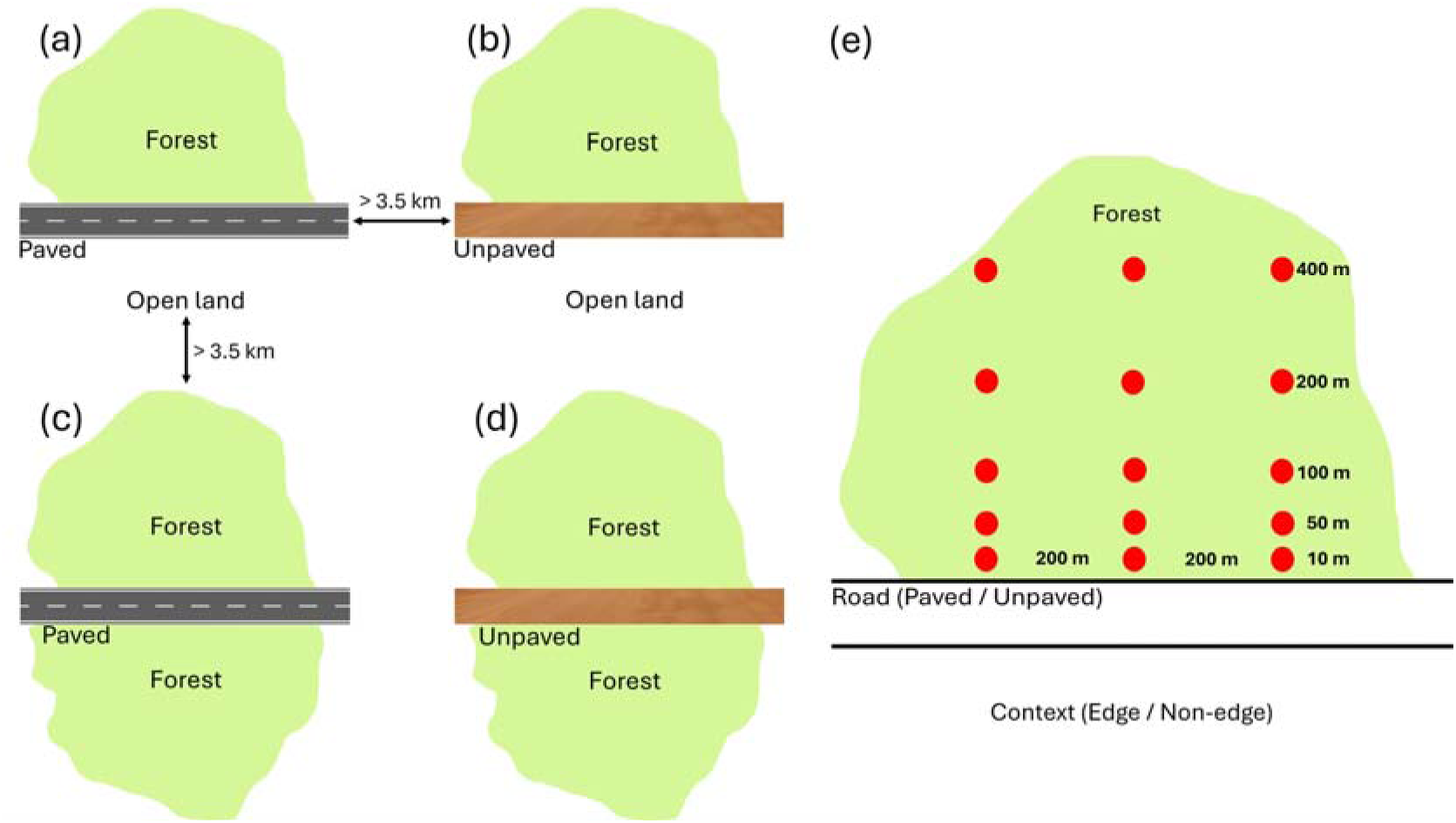
Schematic representation of the sampling layout used to test how rodent-mediated acorn dispersal varies with distance from roads. Road sections were classified by road type (paved vs unpaved) and road–forest context (edge vs non-edge) (a–d). Within each section, seed stations (red circles) were established along a distance gradient into the forest at five target distances from the road edge (10, 50, 100, 200, and 400 m) (e).

### Assessing rodent-mediated seed dispersal

We tracked 3600 labeled holm oak acorns over two consecutive years, from September 2023 to March 2024 and from November 2024 to February 2025. One week before tracking, ripe acorns were collected from 3–4 holm oaks per site. We collected acorns on the tree that fell very easily from the cupules or recently fallen on the ground with a homogeneous brown color and no signs of desiccation (Vaz et al., 2024). Acorns were then mixed and floated in water to remove weevil-infested ones. Mean pre-labeling acorn weight was 7.12 g (± 0.03 SE, range = 2.95–16.35 g). Acorns were labeled by drilling a 1-mm hole (avoiding the embryo) and inserting a 3-cm wire (0.5 mm diameter) attached to a 14.5 cm label (Chen et al., 2019b). Labeling added 3–18% to acorn weight. This method, which does not alter dispersal patterns, ensures high recovery rates (Xiao et al., 2006).

At each site, we established five distances from the road (10, 50, 100, 200 and 400 m) within the forest on one side of the road; when only one side had forest, sampling occurred there (Figure 2B). Each distance included 3 seed stations spaced ∼200 m apart, totaling 15 stations per site (180 supply stations across the 12 sites). To isolate the effect of rodents from other potential consumers, each station was protected with a 32 × 25 × 12 cm wire cage (1.4 cm mesh), containing 10 labeled acorns per year. To allow rodents to the interior, each cage had two 5 × 5 cm entrances and was secured to prevent displacement (Perea et al., 2011a, Vaz et al., 2024). When an oak tree was available (81% of stations), seed stations were placed beneath the tree canopy and, in most cases, positioned about 40 cm from a shrub (usually *Cistus* or *Ulex spp*.). To standardize accessibility, vegetation inside cages was cleared, leaving acorns exposed.

Acorn dispersal was tracked within a minimum 50 m radius around supply stations, expanding by 50 m when a dispersed acorn was found (Craveiro et al., 2025; Vaz et al., 2024). Monitoring followed a decreasing frequency: every 2 days up to day 8, then twice a week for two weeks, weekly for two weeks, biweekly for one month, monthly for two months, and finally bimonthly for four months. Each acorn’s new location was recorded. Depredated acorns (embryo damaged) and those missing (i.e., not found in subsequent searches) were assigned their last recorded dispersal distance. Acorns predated inside supply stations were excluded from dispersal analyses. From week one, non-viable acorns (shriveled, yellowed, hollow) and those predated by other consumers (e.g., wild boars) were removed (Vaz et al., 2024).

Dispersal distance from supply station was measured using a laser meter (1-mm precision). To assess the effect of acorn arrival microhabitat (González-Rodríguez & Villar, 2012; Perea et al., 2011b; Vaz et al., 2024), deposition sites were classified as open areas or under shrubs. Within a 50 m radius of each station, we quantified tree density (trees/ha) by counting trees and visually estimated shrub and grass cover (%) (Galantinho et al., 2020).

To estimate natural acorn availability per year, we counted acorns once beneath and in the crown of the three nearest oak trees to each station (Vaz et al., 2024). Counts were conducted at four cardinal points under the canopy, facing the trunk, with 30 seconds per point. The sum per tree was calculated separately for beneath-canopy and crown samples, and each was averaged across the three trees to obtain station-level estimates. A single observer conducted all counts in the first supplying week for each site.

To examine rodent density effects on acorn dispersal per year, we conducted a 4-night live trapping session at acorn supply stations (Niu et al., 2021; Perea et al., 2011c) at the end of each sampling year (January–March 2024 and January–February 2025). Three Sherman traps were placed at increasing distances from the station (0, 10–20, and 30–50 m) in favorable microsites to maximize captures and improve abundance estimates (under shrubs *Cistus spp*., *Ulex spp.* or tall grass; Fedriani, 2005). Traps were baited with peanut butter on toast (4 × 4 cm), and cotton wool was provided for insulation. Captured rodents were marked with a small lateral fur cut (Hoffmann et al., 2010; Jumeau et al., 2017) and released at capture sites. Procedures complied with Directive 2010/63/EU on animal use in research.

### Analyses – Distance and number of dispersals

To assess the effects of distance to road, road type, road–forest context, year, and covariates (Table 1) on (i) acorn dispersal distance and (ii) the number of dispersals (i.e., number of acorns moved per supply station per year), we fitted two generalized linear mixed models (GLMMs). Because station placement was constrained by local microsite conditions (e.g., proximity to oak canopies and shrubs), stations could not always be placed exactly at the target distances (10, 50, 100, 200, and 400 m). We therefore modelled distance to road using each station’s measured distance as a continuous variable, which captures small placement deviations that are meaningful at the short spatial scale of rodent-mediated dispersal. Correlation analyses indicated collinearity between acorn availability beneath trees and in tree crowns (Spearman’s *r_s_*= 0.71), and between shrub and grass cover (*r_s_* = −0.95). We retained acorn availability beneath trees because these acorns compete with supplied acorns for rodent removal, and shrub cover because it strongly influences rodent-mediated dispersal (Craveiro et al., 2025; Vaz et al., 2024; Yu et al., 2022). To account for potential distance-to-road variation in rodent density (Niu et al., 2021; Ruiz-Capillas et al., 2013), we estimated rodent density separately for each site, year, and target distance class (10, 50, 100, 200, and 400 m) using capture–mark–recapture models in ‘Rcapture’ R package (v1.4-4; Baillargeon & Rivest, 2009) under a closed-population assumption over the 4-night session. We used the bias-corrected closed-population estimator (“closed.bc”). Each station was assigned the density estimate for its distance class.

**Table 1.** Names, definitions, and ranges of variables considered in the analyses of rodent-mediated acorn dispersal distance and number of dispersal events.

| Variable | Description | Range |
| --- | --- | --- |
| Road-forest context | Position of the road relative to the forest | edge, non-edge |
| Road type | Road surface type | paved, unpaved |
| Distance to road | Measured distance (m) from each seed station to the road. Stations placed near target distances (10, 50, 100, 200, and 400 m) |  |
| Year | Sampling year | 1<sup>st</sup> year: Sep 2023–Mar 2024;<br>2<sup>nd</sup> year: Nov 2024–Feb 2025 |
| Acorn weight | Weight (g) | 2.95–16.35 |
| Rodent density | Estimated rodent density (individuals/ha) for each site × year × target distance class (10, 50, 100, 200, 400 m) | 0.00–8.59 |
| Acorn availability | Mean number of acorns on the ground per station per year, estimated from visual counts at the three nearest oaks | 0.00–303.33 |
| Arrival microhabitat | Deposition (arrival) microhabitat of each dispersed acorn | under shrub, open |
| Shrub cover | Visually estimated shrub cover (%) within a 50 m radius of each supply station | 0–86 |
| Grass cover | Visually estimated grass cover (%) within a 50 m radius of each supply station | 6–100 |
| Tree density | Tree density (trees/ha) within a 50 m radius of each supply station | 6.33–335.44 |

For dispersal distances, we fitted a zero-inflated Gamma GLMM with a log link, using acorn dispersal distance from the supply station (m) as the response; zeros indicated no movement, whereas positive values corresponded to the last recorded distance from the station. For the number of dispersals (number of acorns moved per station per year), we fitted a negative binomial type 1 GLMM with a log link to account for overdispersion in count data. Random effects reflected the sampling design: the dispersal-distance model included station nested within site (1 | site/station), whereas the number-of-dispersals model included site as a random intercept (1 | site). In both models, year was included as a fixed effect and was also specified in the zero-inflation and dispersion components (ziformula = ∼ year; dispformula = ∼ year) to allow the frequency of zeros and residual variance to vary between years.

Final models were obtained by fitting a priori global GLMMs for each response and simplifying them to a parsimonious set of predictors. Global models included our hypothesized moderators of distance-to-road effects (interactions of distance to road with road–forest context, road type, and year) alongside candidate covariates describing local habitat and acorn availability (Table 1). We then performed backward elimination, sequentially removing non-significant terms based on likelihood-ratio tests (LRTs) for nested models (Zuur et al., 2009). The final dispersal-distance model retained the road–forest context × distance to road and year × distance to road interactions, as well as arrival microhabitat (under shrub vs open), acorn availability beneath trees, and tree density, with station nested within site as random effects. The final model for the number of dispersals retained road type, distance to road, shrub cover, and year, with site as a random intercept.

Models were implemented in the ‘glmmTMB’ R package (v1.1.7; Brooks et al., 2017). Model adequacy was assessed using the ‘DHARMa’ R package (v0.4.6; Hartig, 2022), which generates residual diagnostics, including QQ-plots, dispersion tests, and residuals vs. predicted values plots. We assessed spatial independence of station□level mean residuals (residuals averaged per station) using a Monte Carlo Moran’s I test (999 permutations; spdep::moran.mc). The observed Moran’s I was –0.006 (two□sided *p* = 0.110), indicating no significant spatial autocorrelation of residuals. Analyses were conducted in R version 4.1.1 (R Core Team, 2021).

## RESULTS

Across both years, rodents dispersed 1219 of the 3600 supplied acorns, with a mean dispersal distance of 2.27 m (Table 2). Dispersal distances were generally greater at stations farther from roads (100–400 m) than at stations closer to roads (10–50 m), whereas the number of dispersal events tended to be higher near roads than farther into the forest. No road crossings were detected.

**Table 2.**
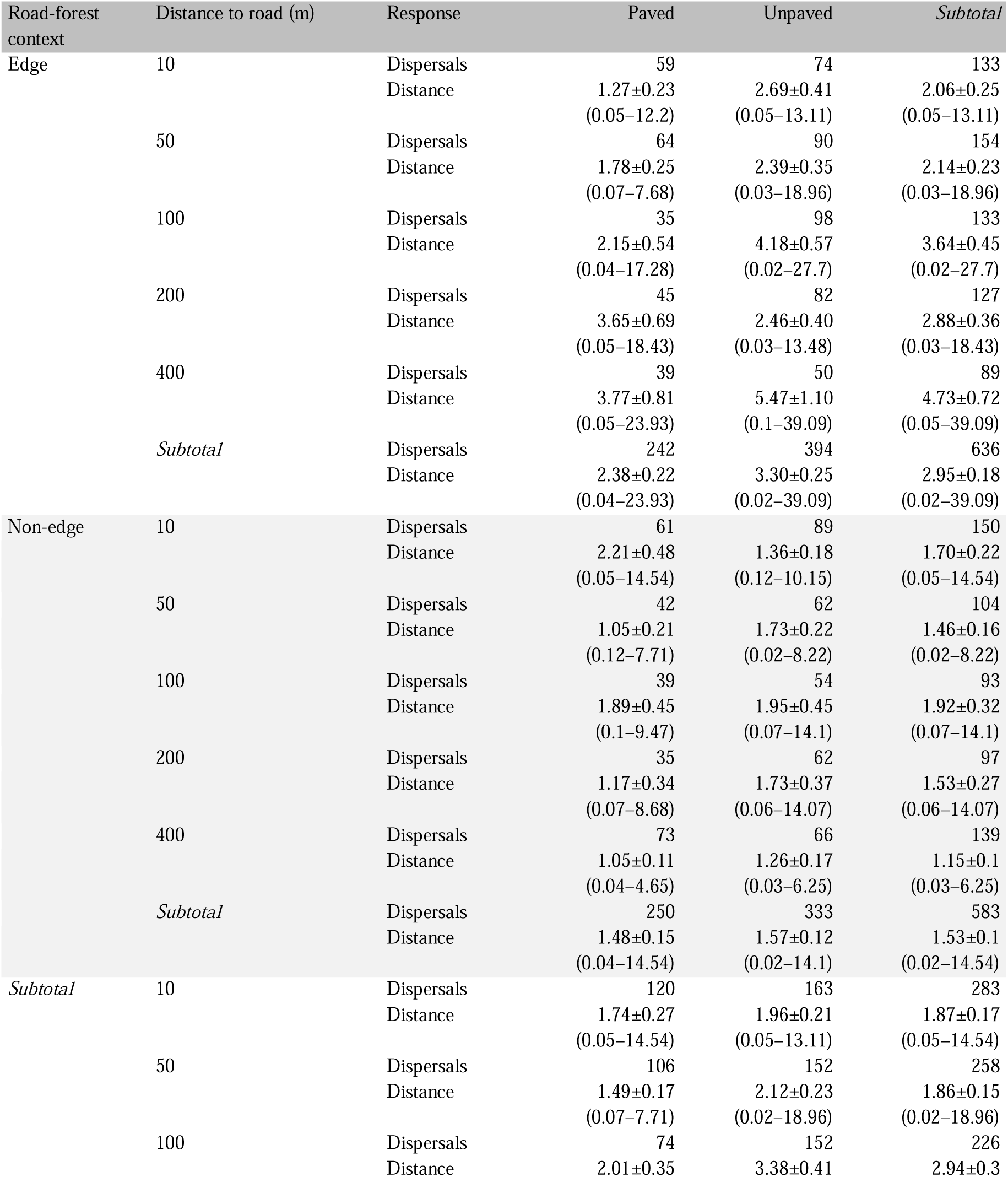
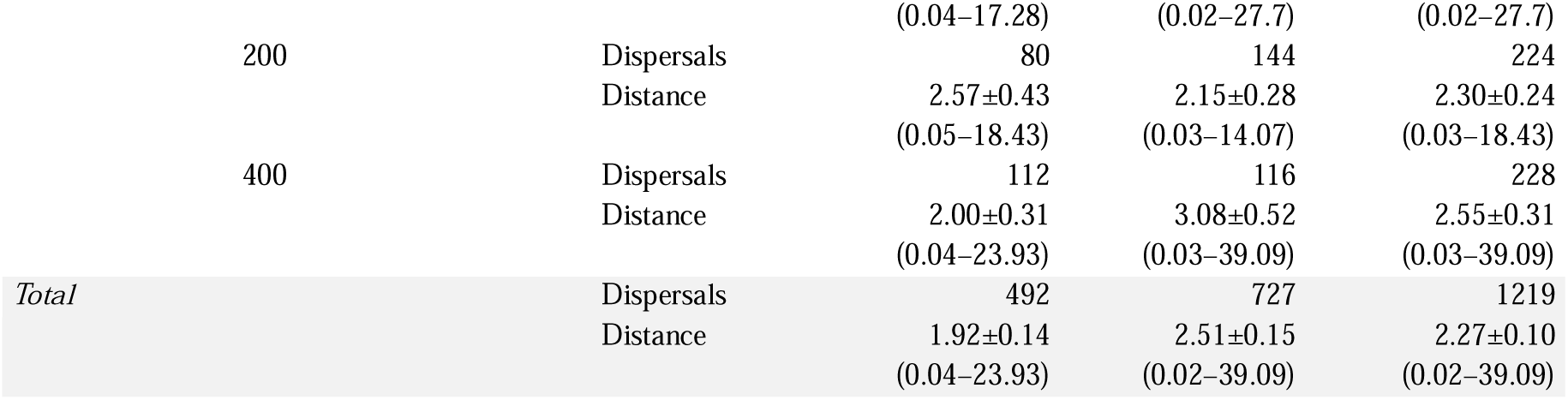
Observed seed dispersal metrics for rodents across edge and non-edge road-forest contexts with paved and unpaved roads, and varying distances to road (m). Shown are the total number of dispersals and dispersal distances (m) (mean ± SE, range).

| Road-forest context | Distance to road (m) | Response | Paved | Unpaved | Subtotal |
| --- | --- | --- | --- | --- | --- |
| Edge | 10 | Dispersals | 59 | 74 | 133 |
| | | Distance | 1.27±0.23<br>(0.05–12.2) | 2.69±0.41<br>(0.05–13.11) | 2.06±0.25<br>(0.05–13.11) |
|  | 50 | Dispersals | 64 | 90 | 154 |
| | | Distance | 1.78±0.25<br>(0.07–7.68) | 2.39±0.35<br>(0.03–18.96) | 2.14±0.23<br>(0.03–18.96) |
|  | 100 | Dispersals | 35 | 98 | 133 |
| | | Distance | 2.15±0.54<br>(0.04–17.28) | 4.18±0.57<br>(0.02–27.7) | 3.64±0.45<br>(0.02–27.7) |
|  | 200 | Dispersals | 45 | 82 | 127 |
| | | Distance | 3.65±0.69<br>(0.05–18.43) | 2.46±0.40<br>(0.03–13.48) | 2.88±0.36<br>(0.03–18.43) |
|  | 400 | Dispersals | 39 | 50 | 89 |
| | | Distance | 3.77±0.81<br>(0.05–23.93) | 5.47±1.10<br>(0.1–39.09) | 4.73±0.72<br>(0.05–39.09) |
|  | *Subtotal* | Dispersals | 242 | 394 | 636 |
| | | Distance | 2.38±0.22<br>(0.04–23.93) | 3.30±0.25<br>(0.02–39.09) | 2.95±0.18<br>(0.02–39.09) |
| Non-edge | 10 | Dispersals | 61 | 89 | 150 |
| | | Distance | 2.21±0.48<br>(0.05–14.54) | 1.36±0.18<br>(0.12–10.15) | 1.70±0.22<br>(0.05–14.54) |
|  | 50 | Dispersals | 42 | 62 | 104 |
| | | Distance | 1.05±0.21<br>(0.12–7.71) | 1.73±0.22<br>(0.02–8.22) | 1.46±0.16<br>(0.02–8.22) |
|  | 100 | Dispersals | 39 | 54 | 93 |
| | | Distance | 1.89±0.45<br>(0.1–9.47) | 1.95±0.45<br>(0.07–14.1) | 1.92±0.32<br>(0.07–14.1) |
|  | 200 | Dispersals | 35 | 62 | 97 |
| | | Distance | 1.17±0.34<br>(0.07–8.68) | 1.73±0.37<br>(0.06–14.07) | 1.53±0.27<br>(0.06–14.07) |
|  | 400 | Dispersals | 73 | 66 | 139 |
| | | Distance | 1.05±0.11<br>(0.04–4.65) | 1.26±0.17<br>(0.03–6.25) | 1.15±0.1<br>(0.03–6.25) |
|  | *Subtotal* | Dispersals | 250 | 333 | 583 |
| | | Distance | 1.48±0.15<br>(0.04–14.54) | 1.57±0.12<br>(0.02–14.1) | 1.53±0.1<br>(0.02–14.54) |
| *Subtotal* | 10 | Dispersals | 120 | 163 | 283 |
| | | Distance | 1.74±0.27<br>(0.05–14.54) | 1.96±0.21<br>(0.05–13.11) | 1.87±0.17<br>(0.05–14.54) |
|  | 50 | Dispersals | 106 | 152 | 258 |
| | | Distance | 1.49±0.17<br>(0.07–7.71) | 2.12±0.23<br>(0.02–18.96) | 1.86±0.15<br>(0.02–18.96) |
|  | 100 | Dispersals | 74 | 152 | 226 |
| | | Distance | 2.01±0.35 | 3.38±0.41 | 2.94±0.3 |

|  |  |  |  |  |
| --- | --- | --- | --- | --- |
|  |  | (0.04–17.28) | (0.02–27.7) | (0.02–27.7) |
| 200 | Dispersals | 80 | 144 | 224 |
|  | Distance | 2.57±0.43 | 2.15±0.28 | 2.30±0.24 |
|  |  | (0.05–18.43) | (0.03–14.07) | (0.03–18.43) |
| 400 | Dispersals | 112 | 116 | 228 |
|  | Distance | 2.00±0.31 | 3.08±0.52 | 2.55±0.31 |
|  |  | (0.04–23.93) | (0.03–39.09) | (0.03–39.09) |
| *Total* | Dispersals | 492 | 727 | 1219 |
|  | Distance | 1.92±0.14 | 2.51±0.15 | 2.27±0.10 |
|  |  | (0.04–23.93) | (0.02–39.09) | (0.02–39.09) |

Importantly, natural acorn availability was 64% lower in second year than first year (mean ± SE: 30.8 ± 3.3 vs 85.0 ± 5.1 acorns per tree), indicating a poor mast year. By the end of the monitoring period, 81% of dispersed acorns had been consumed, 33% of supplied acorns were missing, and 22% were predated by livestock.

Live trapping indicated that Algerian mice (63% of captures) and wood mice (37%) mediated most dispersal activity, with a combined estimated density of 2.84 individuals ha□¹. Rodent density tended to decrease with increasing distance from roads (3.41 vs. 2.37 individuals ha□¹ at 10 and 400 m, respectively), while intermediate distances showed slightly higher values (2.66–2.94 individuals ha□¹ at 50–200 m). Density was 2.9 times higher in the second year than in the first year.

### 3.1 Effects on distance and number of dispersals

The final mixed model for acorn dispersal distance (Table 3) indicated that distance-to-road effects varied with road–forest context and year (Figure 3). In non-edge contexts (roads traversing forest), dispersal distance showed little evidence of change along the distance-to-road gradient (Figure 3a). By contrast, in edge contexts, predicted distances were 2× higher at 400 m (3.08 m) than at 10 m (1.55 m). Between years, distance-to-road effects were weak in year 1 but became evident in year 2 (poor mast year), with predicted distances at 400 m reaching 2× those at 10 m (Figure 3b). We found no evidence that road type affected dispersal distance, nor that it modified distance-to-road effects.

**Figure 3.**
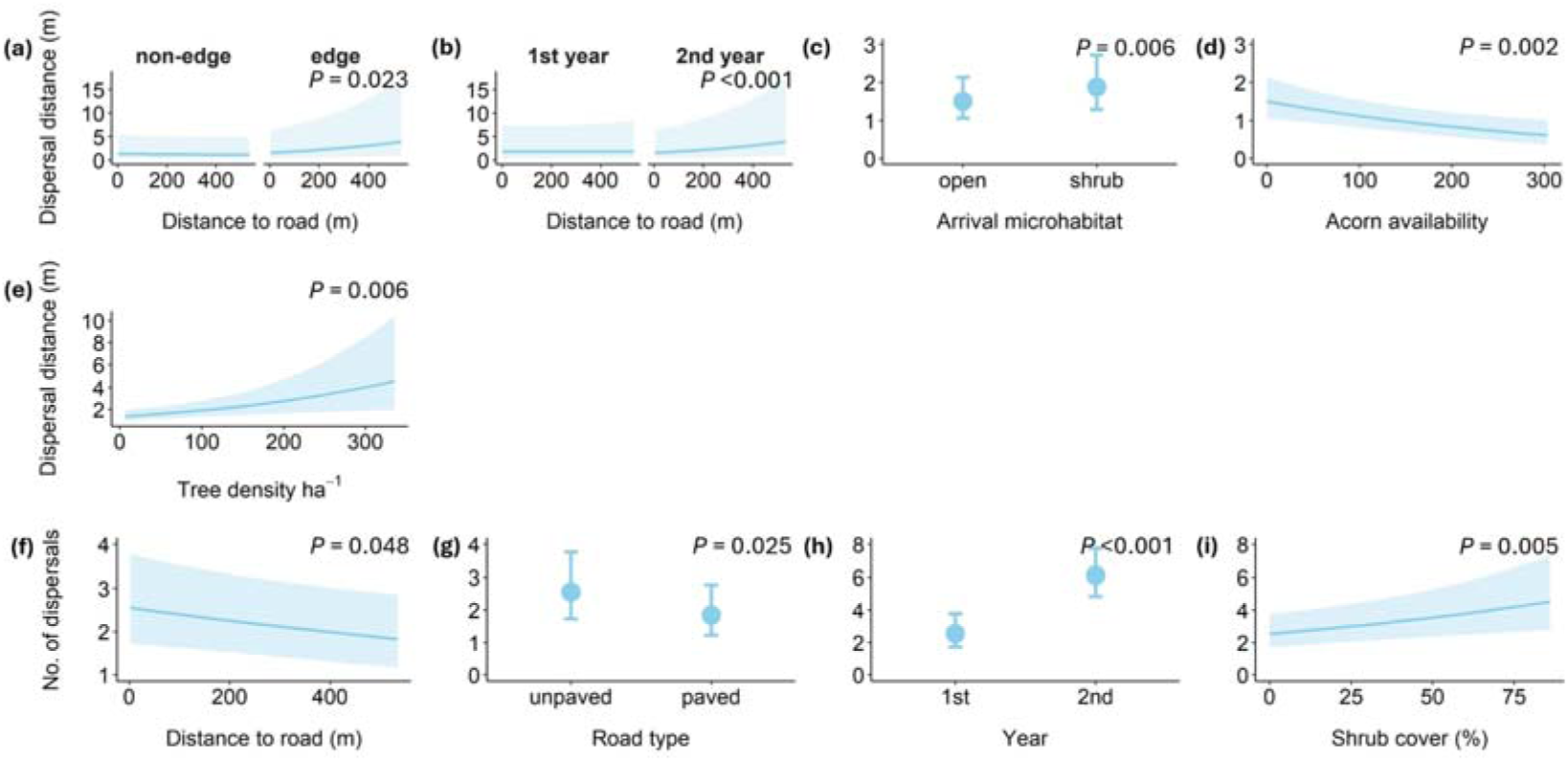
Mean fitted values (±95% CI) from the optimal mixed-effects models predicting (a–e) acorn dispersal distance and (f–i) number of acorn dispersals mediated by rodents.

**Table 3.**
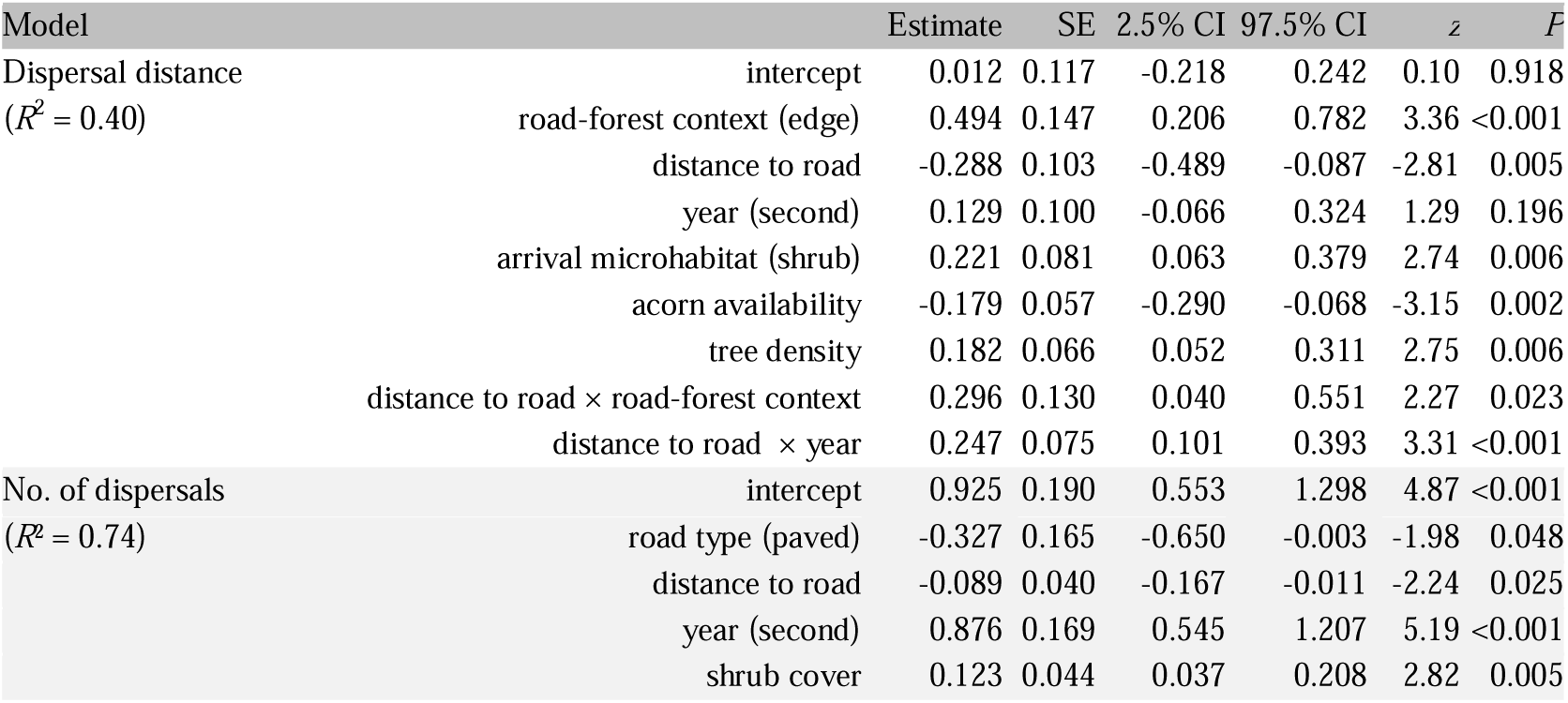
Parameter estimates (estimate ± SE), 95% confidence intervals, *z*-values, and *P*-values for fixed effects in the final GLMMs predicting (i) acorn dispersal distance and (ii) number of dispersal events. Reference levels: road–forest context = non-edge; road type = unpaved; year = 1^st^ year; arrival microhabitat = open. *R*² values are marginal, based on Nakagawa’s *R*² framework.

Dispersal distances also depended on arrival microhabitat (Figure 3c): acorns deposited under shrubs were dispersed farther than those deposited in open microsites (1.87 m vs 1.50 m; 25% increase). Dispersal distances decreased with increasing natural acorn availability (Figure 3d), dropping by 45% as availability increased from 100 to 300 acorns (from 1.12 m to 0.68 m). Finally, dispersal distances increased with tree density (Figure 3e), rising 2.1-fold as density increased from 100 to 300 trees ha□¹ (from 1.92 m to 4.03 m).

In the model for the number of dispersal events (Table 3), dispersal frequency decreased with distance from the road (Figure 3f), with 30% more dispersal events at 10 m than at 400 m, consistent with our expectations. As hypothesized, road type also influenced dispersal frequency (Figure 3g), with seed stations along unpaved roads sustaining 39% more dispersal events than those along paved roads. The model found no support for an effect of road–forest context on dispersal frequency, nor for an interaction with distance from the road.

Dispersal frequency also differed between years (Figure 3h), increasing 2.4-fold from year 1 to year 2 (poor mast year). Finally, dispersal frequency increased with shrub cover (Figure 3i), rising by ∼40% from 25% to 75% shrub cover.

## DISCUSSION

Road proximity reshaped rodent-mediated acorn dispersal in two distinct ways: it increased dispersal activity near roads but did not translate into longer dispersal distances there, and it did not promote cross-road seed movement. Instead, dispersal distances depended on road–forest context and interannual variation, being most constrained near roads in edge contexts and during the low-acorn year, while dispersal events were most frequent near roads and along unpaved roads. Together, these results indicate that roadside verges can act as high-activity zones for scatter-hoarding rodents, but their contribution to regeneration depends on whether dispersed acorns are deposited in structurally safe microsites (e.g., shrubs), highlighting opportunities for verge and understory management to enhance deposition in recruitment-favorable microsites and mitigate near-road constraints.

High dispersal activity near roads did not translate into longer dispersal distances. Dispersal events were most frequent near roads, consistent with the idea that road verges can concentrate rodent activity and seed handling, especially in managed landscapes where verges may provide predictable resources or refuge structure (Ascensão et al., 2012; Galantinho et al., 2020; Ruiz-Capillas et al., 2013). Yet near-road conditions did not yield longer dispersal distances; rather, dispersal distances were constrained near roads in edge contexts and varied with year. This decoupling of dispersal frequency and distance indicates that roads can increase removal while shortening dispersal distances, thereby altering dispersal effectiveness in ways that cannot be inferred from removal rates alone (Cui et al., 2018; Niu et al., 2018; Chen et al., 2019a, 2019b).

Road–forest context governed the road-distance signal in dispersal distances. In non-edge contexts (roads traversing forest), dispersal distances showed little evidence of change across the road-distance gradient, whereas in edge contexts dispersal distances increased farther into forest interiors. This context dependence aligns with evidence that edge environments reshape rodent foraging and caching behavior by altering cover, microclimate, and perceived predation risk (López-Barrera et al., 2007; Mazzamuto et al., 2018; Morán-López et al., 2016). Roads bordered by open areas may intensify forest-side edge effects because verges are often more disturbed and structurally simplified than roads embedded within forest on both sides (Craveiro et al., 2025; Galantinho et al., 2017), creating sharp structural contrasts that can constrain dispersal distances near the road edge while allowing longer dispersal distances farther from roads within the forest. Overall, these patterns are inconsistent with a simple linear distance–decay relationship and instead reflect interactions between road disturbance and structural habitat heterogeneity (Mazzamuto et al., 2018; Morán-López et al., 2016).

Roads also remained functionally impermeable to acorn movement across the road surface even where rodent activity was elevated near roads. No acorns crossed the road, consistent with strong barrier effects of roads for rodents carrying seeds and the idea that exposure on road surfaces discourages crossings with acorns (Craveiro et al., 2025; Niu et al., 2021). Consequently, even where roadsides support frequent seed handling, the benefits are likely confined to the forest side adjacent to roads rather than extending across road surfaces. Although we tracked acorn fate over a period long enough to detect seedling emergence, we observed no seedlings. Thus, implications for regeneration should be interpreted through the well-established role of scatter-hoarding in recruitment, while acknowledging that establishment ultimately depends on post-dispersal processes and microsite conditions (Arousa et al., 2015; Perea et al., 2011a; Vaz et al., 2024).

Road type added nuance by affecting dispersal frequency but not dispersal distance. Unpaved roads sustained more dispersal events than paved roads, whereas dispersal distances showed no road-type effect and no modification of the distance-to-road gradient. This contrasts with contexts where paved road verges have been associated with longer movements or different movement orientations (Craveiro et al., 2025), reinforcing that “road type” likely captures correlated features such as traffic, verge management, and verge structure whose effects depend on local conditions. More broadly, our pattern of higher near-road dispersal activity coupled with shorter dispersal distances near roads is compatible with verge-driven concentration of rodent activity and constraints on longer movements in the road vicinity (Chen et al., 2019a, 2019b; Cui et al., 2018; Niu et al., 2018). Higher local rodent densities may further shorten transport distances via interference competition and cache pilferage risk (Cao et al., 2018; Dally et al., 2006).

Interannual variation suggests that mast context can modulate how road proximity shapes dispersal distances. Dispersal distances showed a year × distance-to-road interaction, becoming more evident in the low-acorn year, while dispersal frequency increased overall in that year. Annual fluctuations in acorn availability are known to shift rodent foraging decisions and the balance between consumption and caching, with implications for both how many seeds are handled and how far they are moved (Puerta-Piñero et al., 2010; Vaz et al., 2024; Wang et al., 2017). In poor mast years, when each acorn is effectively more valuable, rodents may become more risk-averse while transporting seeds, especially in exposed microsites near roads; such exposure can be reinforced where verge environments have lower canopy cover than adjacent forest, increasing perceived predation risk and constraining longer movements in the most open conditions (Delgado et al., 2007; Yu et al., 2022). Taken together, these patterns indicate that assessing dispersal services in roaded landscapes may require accounting for interannual variation in acorn availability, particularly when the goal is to identify when and where roads constrain dispersal distances within adjacent forests (Puerta-Piñero et al., 2010; Vaz et al., 2024; Wang et al., 2017).

Fine-scale habitat structure provides a mechanistic bridge between verge-associated activity and dispersal outcomes. Dispersal distances were longer when acorns were deposited under shrubs than in open microsites, and shrub cover increased dispersal frequency. These results align with evidence that wood mice and Algerian mice preferentially use shrub-dense areas for movement and caching because shrubs provide cover and reduce perceived predation risk (Galantinho et al., 2022; Morán-López et al., 2015; Perea et al., 2011c; Rosalino et al., 2011; Sun et al., 2013; Vaz et al., 2024). Work in other systems suggests threshold-like relationships between shrub cover and dispersal/caching activity (e.g., ∼65% shrub cover; Morán-López et al., 2016), reinforcing shrub structure as a key determinant of dispersal dynamics. Shrubs can also act as nurse microsites that enhance survival and establishment under Mediterranean climatic stress (Andivia et al., 2017; Perea et al., 2016), although they may simultaneously increase seed handling and predation in some circumstances (Perea et al., 2011a). Conversely, open microsites can sometimes favor growth via higher light availability and reduced competition, but may increase drought stress and mortality in hot, dry summers (Arousa et al., 2015; Yu et al., 2022). Thus, our results support the practical value of maintaining a mosaic of shrub patches and open areas that can provide both sheltered dispersal pathways/caching sites and microsites suitable for subsequent establishment (Andivia et al., 2017; Arousa et al., 2015; Perea et al., 2011a, 2016).

Stand structure and local resource availability further shaped dispersal distances, indicating multiple interacting constraints on dispersal services. Dispersal distances increased with tree density, consistent with the idea that greater canopy cover and structural complexity can reduce perceived predation risk and facilitate movement within safer environments (Rosalino et al., 2011; Wang et al., 2025). In contrast, greater natural acorn availability reduced dispersal distances, consistent with the expectation that abundant local resources reduce incentives for long-distance transport and favor local consumption or short-distance caching (Niu et al., 2021; Vaz et al., 2024). These patterns suggest that road proximity operates within a broader template of habitat structure and seed availability, and that management actions aimed at strengthening dispersal services should consider how structural conditions and mast context jointly shape rodent decisions over fine spatial scales.

### Management implications

Our findings provide actionable guidance for enhancing oak regeneration in roaded Mediterranean woodlands by identifying where and how verge-associated rodent activity can be translated into dispersal outcomes more likely to support recruitment. Road verges concentrated seed handling—especially near roads and along unpaved roads—yet this elevated activity did not produce longer dispersal distances near roads and did not generate cross-road seed movement. Management should therefore not assume that high activity at verges automatically promotes regeneration, nor that rodents will supply functional connectivity across road surfaces. Instead, the management goal should be to increase the likelihood that acorns handled near verges are deposited into recruitment-favorable microsites.

Our results identify the verge–forest interface as a critical leverage point, particularly in edge contexts where dispersal distances were most constrained near roads but increased farther into the forest. In landscapes where roads border open habitats, abrupt transitions likely increase exposure and perceived risk near the road, limiting transport distances. Actions that maintain or restore structural complexity at the boundary—rather than simplifying it through uniform clearing—are therefore likely to mitigate near-road constraints. In our study, shrub-associated deposition increased dispersal distances and shrub cover increased dispersal frequency, indicating that structural cover is central to both the occurrence of dispersal events and the delivery of acorns into sheltered microsites. Where compatible with road safety and fire prevention, verge maintenance that avoids uniform clearing and instead retains shrub patches and heterogeneous understory structure should help sustain rodent activity while increasing the probability that moved acorns end up in safe deposition sites. Because tree density also increased dispersal distances, interventions that preserve or restore stand structure adjacent to roads are likely to further facilitate these outcomes.

Finally, management should be responsive to mast context. Distance-to-road effects on dispersal distances became most evident during the low-acorn year, suggesting that poor mast years may be periods when vegetation structure and edge conditions exert stronger control over how far rodents transport acorns relative to roads. In applied terms, this implies that management actions targeting structural cover at verges and into forest interiors may be particularly valuable—and their effects more detectable—during years of low natural acorn availability. Our results support a straightforward applied message for roaded oak landscapes: maintaining heterogeneous structural cover at road verges and in adjacent understory can harness verge-associated rodent activity to strengthen delivery of acorns into recruitment-favorable microsites, even where cross-road seed movement is effectively absent.

## CRediT authorship contribution statement

**João Craveiro**: Data curation, Formal analysis, Investigation, Visualization, Writing – original draft. **Miguel N. Bugalho**: Supervision, Writing – review & editing. **Pedro G. Vaz**: Conceptualization, Supervision, Resources, Writing – review & editing.

## Acknowledgments

JC was supported by the Portuguese Science and Technology Foundation (FCT) through an individual grant (2021.05551.BD) and institutional funding to CEABN-InBIO (UIDB/50027/2025) and InBIO (DOI 10.54499/LA/P/0048/2020). JC and PGV were funded by FCT through CE3C (DOI 10.54499/UIDB/00329/2020) and CHANGE (DOI 10.54499/LA/P/0121/2020). PGV received funding from the AdaptForGrazing project (PRR-C05-i03-I-000035-LA4.3/4.4/4.6/4.7), supported by the EU (RRF) via IFAP (PRR).

